# Gene editing in the nematode parasite *Nippostrongylus brasiliensis* using extracellular vesicles to deliver active Cas9/guide RNA complexes

**DOI:** 10.1101/2022.08.30.505940

**Authors:** Jana Hagen, Subhanita Ghosh, Peter Sarkies, Murray E. Selkirk

## Abstract

Despite recent advances, animal-parasitic nematodes have thus far been largely refractory to genetic manipulation. We describe here a new approach providing proof of principle that CRISPR/Cas9-mediated gene editing of parasitic nematodes is achievable using vesicular stomatitis virus glycoprotein-pseudotyped extracellular vesicles (EV) for the delivery of Cas9-synthetic guide RNA (RNP) complexes. We demonstrate that EV-delivered RNPs can be used to disrupt a secreted DNase II in *Nippostrogylus brasiliensis.* Introduction of a repair template encoding multiple stop codons led to measurable reduction in expression of the targeted gene. Altered transcripts corresponding to the edited locus were detected by RT-PCR, demonstrating that vesicles can access cells of tissues actively expressing the gene of interest. These data provide evidence that this technique can be employed for targeted gene editing in *N. brasiliensis,* making this species genetically tractable for the first time and providing a new platform for genetic analysis of parasitic nematodes.

**Author Summary:** Parasitic nematodes have a complex life cycle involving passage through a host organism, which makes them very difficult to manipulate genetically. Recently, a method for deleting, changing or replacing genes (gene editing) has been developed in other organisms which has revolutionised our ability to understand fine details of how these organisms work. It has generally not been possible to adapt this method to parasitic nematodes because delivery of the components is difficult, and this has proved to be a bottleneck in understanding how parasites develop, survive and interact with their host. We show here that the components for gene editing can be introduced into a widely used laboratory model of intestinal nematode infection by encapsulation in membrane-bound vesicles which have been modified to carry a protein which facilitates fusion of the vesicles with parasite cells and delivery of the contents. This resulted in accurate editing of a specific gene by deletion and repair, such that the amount of functional protein produced from that gene was reduced. This system should be applicable to all nematode species, and will facilitate understanding of their complex biology, in addition to defining new targets for control of infection.

## Introduction

Over a quarter of the world’s population are estimated to be infected with soil-transmitted helminths, representing a severe burden of disease and disability [1]. Additionally, gastrointestinal nematodes are responsible for major economic losses to the livestock industry, with rising multi-drug resistance to the major classes of anthelmintics [2]. A major bottleneck to identifying molecules that might serve as new drug targets or vaccine candidates is the genetic intractability of most parasitic nematodes, which limits screening proteins for biological properties and essential functions.

The delivery of foreign DNA or RNA has been identified as a limiting factor for gene silencing and transgenesis in parasitic nematodes [3,4]. Recently, we showed that *Nippostrongylus brasiliensis* could be transduced with vesicular stomatitis virus glycoprotein (VSV-G)-pseudotyped lentiviral particles, with entry most likely mediated by binding of VSV-G to low-density lipoprotein receptor-related (LRP) proteins [5,6]. However, the delivered expression cassette was subjected to gene silencing during worm development, such that further optimisation of the system is required in order to achieve robust and reliable transgene expression [5].

Clustered Regularly Interspaced Short Palindromic Repeat (CRISPR)/Cas9 mediated gene editing is a powerful tool which facilitates permanent modifications in genomic DNA, and this has been successfully applied to a range of helminth species for the generation of gene knockout and knockin mutants [7–9]. However, delivery of expression cassettes or Cas9/synthetic guide (sg)RNA ribonucleoprotein (RNP) complexes to parasitic nematodes has proven problematic. Injection into the syncytial gonad of adult female *Strongyloides* has been successful [7], as has lipofection of *Brugia malayi* with micelles containing RNP complexes [10]. These approaches were possible because of the unique free-living phase of *Strongyloides,* and by development of a culture system for *B. malayi,* which facilitated genetic manipulation of parasites and selection of mutants [11].

Recently, a new approach was established for transient delivery of RNP complexes into mammalian cells using VSV-G-pseudotyped extracellular vesicles, named NanoMEDIC (nano<u>m</u>embrane-derived <u>e</u>xtracellular vesicles for the <u>d</u>el<u>i</u>very of macromolecular <u>c</u>argo) [12]. Proof of principle was demonstrated in several cell types which had proven difficult to transfect such as induced pluripotent stem cells, neurons and myoblasts [12]. We thought that this could be a useful tool to apply to helminths, as lengthy optimisation of the viral delivery system for individual parasite species would be circumvented by production and assembly of RNP complexes in mammalian cells for which optimised expression cassettes and transfection protocols are readily available. Following the success of VSV-G-mediated uptake of lentiviral particles, we investigated whether NanoMEDIC could be utilised in *N. brasiliensis* as a model gastrointestinal nematode.

## Results

### Characterisation of *dnase2* expression in different life stages

To assess susceptibility of CRISPR-mediated gene editing in *N. brasiliensis,* we chose secreted DNase II as a target gene [13], as endonuclease activity in secreted products provides a means for functional analysis of gene expression. While the DNase II has been described to be secreted from infective larvae [14], a profile of expression in different stages has not been determined. Because a shift in temperature to 37°C acts as a cue for exsheathment and initiation of feeding in infective larvae (L3) [15], we first analysed levels of DNase II transcripts and secretion of active enzyme in L3 prior to and during activation by culture at 37°C in the presence of rat serum. Incubation at the elevated temperature led to a sharp increase in DNase II transcripts, peaking at an approximate 1000-fold increase after 2-3 days (Fig 1A). Transcript levels were drastically reduced in adult worms compared with activated L3, but remained approximately 5-fold above those in non-activated L3 (Fig 1A). Western blot analysis of parasite secreted products confirmed that the DNase II was released predominantly by activated L3 (Fig 1B). Given the detection of low levels of DNase II transcripts in non-activated L3, there was a possibility that the protein might be pre-synthesised and stored in secretory vesicles allowing for immediate release upon entering the host. To assess this, we exposed non-activated and activated L3 to plasmid DNA during culture at 37°C. However, non-activated L3 did not secrete sufficient amounts of DNase II for effective degradation of DNA within the first 2 hours of incubation at 37°C, although complete degradation was achieved following overnight incubation (Fig 1C). In contrast, degradation of DNA in the culture supernatant was observed immediately following exposure to previously activated L3.

**Fig 1.**
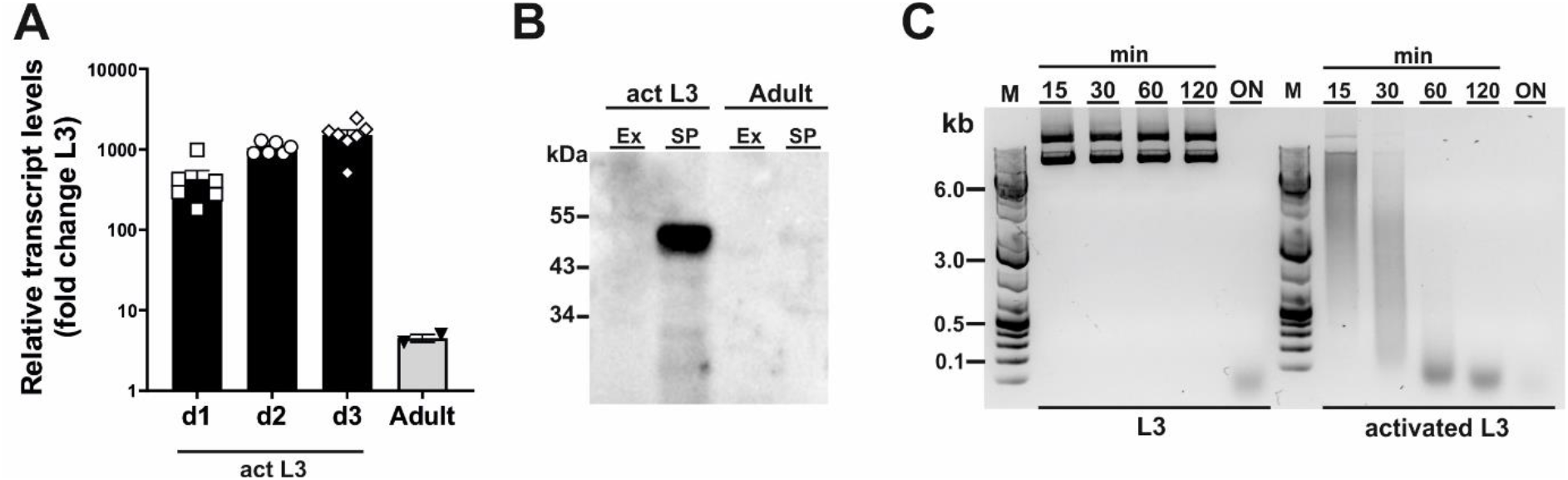
*Dnase2* expression in *Nippostrongylus brasiliensis.* (A) *Dnase2* transcripts are upregulated in L3 following activation at 37°C. Freshly isolated L3 were washed extensively, incubated at 37°C, and collected after 1 to 3 days of in vitro culture. Adult worms were recovered from the intestine of rats at day 8 post-infection. Activated L3 and adult worms were analysed by RT-qPCR for *dnase2* transcript levels relative to that of unactivated L3. (B) DNase II is predominantly secreted by activated L3. Adult worms or activated L3 were incubated for 7 days or 14 days respectively in serum-free medium, culture supernatants collected and concentrated. Subsequently, 5 ug of secreted products (SP) or worm extract (Ex) were analysed for the presence of DNase II by western blot. (C) Non-activated L3 do not readily secrete DNase II. Freshly isolated or activated L3 (200 worms) were incubated in 100 μl of serum-free medium containing 2 μg of plasmid DNA for 15 min to 18 hours (overnight, ON) at 37°C. Supernatants were collected at various time points and assessed for plasmid DNA degradation by resolution of DNA fragments on 1% agarose gels.

### Generation of extracellular vesicles containing DNase II-ribonucleoprotein complexes or a single stranded oligodeoxynucleotide

The genomic DNA sequence of DNase II was identified in NBR_scaffold_0001590 following alignment of the published cDNA sequence (GenBank: M938457) [13] to the genomic *N. brasilensis* database PRJEB511 available on WormBase ParaSite. Nine exons were defined with the two HxK motifs characteristic of DNase II and which together form a single active site [16] encoded in exons 3 and 7 (Fig 2A). Notably, while the corresponding amino acid sequence derived from NBR_scaffold_0001590 and M938457 was mainly conserved, some differences were detected in the cDNA sequences which could affect the prediction of effective sgRNAs (Suppl 1). Expression of NB_Dnase2_1590 was confirmed in our laboratory strain of *N. brasiliensis* following cloning of the cDNA sequence into a yeast expression vector and subsequent sequencing such that sgRNA design was based on NB_Dnase2_1590.

**Fig 2.**
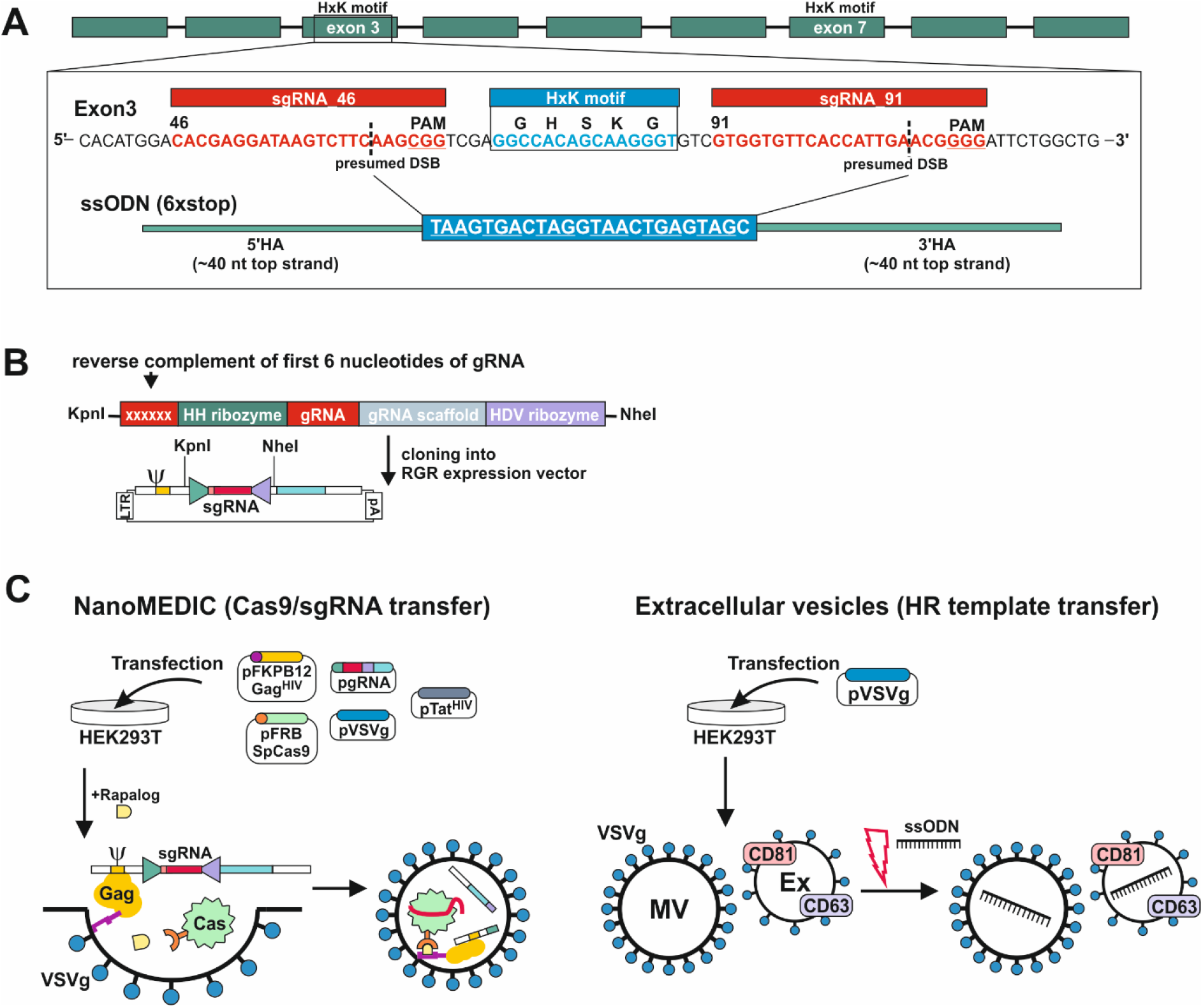
Generation of extracellular vesicles carrying ribonucleoprotein complexes or a homology directed repair template. (A) Gene model of *dnase2* (NBR_1590) showing the position of its nine exons, eight introns and the location of the two HxK motifs. Inset of exon three: nucleotide sequence of the (+) strand indicating location and sequence of gRNA target sites (red), protospacer adjacent motifs (PAM, underlined), the presumed double-stranded breaks (DSB) (dashed line, and the 119-nucleotide sequence of the single-stranded DNA donor template provided for DSB repair by homologous recombination. Homology arms of 45 or 50 nt flank the central 24 nt of a six-stop-codon transgene. (B) Small guide (sg)RNA sequences were cloned between the self-cleaving Hammerhead (HH) and hepatitis delta virus (HDV) ribozymes of the transfer plasmid (ribozyme-guide-ribozyme, RGR) encoding a virus-like mRNA with a packaging signal sequence (Ψ). (C) VSV-G-pseudotyped EVs containing Cas9/gRNA complexes (NanoMEDIC) were produced in HEK293T cells following transfection. The modification of Gag with FKB12 (FK506 binding protein) ensures effective packing of Gag/mRNA complexes following binding to the cell membrane. Addition of a rapalog (rapamycin analogue) ligand mediates heterodimerisation of FRB (rapamycin-binding domain)-modified Cas9 with FKB12-Gag. The sgRNA encoded in virus-like mRNA is liberated in NanoMEDIC following cleavage by two self-cleaving ribozymes and incorporated into Cas9. (D) For the HDR template transfer, EVs were produced by VSV-G-expressing HEK293T cells. Evs consisting of microvesicles (MV) and exosomes (Ex) were then loaded with HDR templates by electroporation.

For transient delivery of Cas9/gRNA complexes, we adapted a recently developed NanoMEDIC approach established in mammalian cells using VSV-G-pseudotyped extracellular vesicles [12]. To predict sgRNA target sequences, the Cas-Designer (RGEN) algorithm was used, which allows for off-target screening against the *N. brasiliensis* genome [17,18]. We chose exon 3 as a target region as it encodes one of the two HxK motifs of functional DNases [16] (Fig 2A). While some gRNAs were predicted that covered the HxK region, these did not achieve a frameshift prediction score over 66. The two highest scoring gRNAs, guide 46 and 91, flanking the HxK motif in exon 3, were cloned between the self-cleaving Hammerhead (HH) and hepatitis delta virus (HDV) ribozymes of the transfer plasmid (ribozyme-guide-ribozyme, RGR) (Fig 2B). If the deletion was insufficient or frameshift not achievable using either guide, then the EV approach would allow for multiplexing to include both guide RNAs deleting the entire motif coding sequence. Corresponding VSV-G-pseudotyped EVs containing RNP complexes (NanoMEDIC) were produced in HEK293T cells following transfection with the RGR and four packaging plasmids encoding Cas9, Gag, VSV-G and Tat (Fig 2C) [12].

Long terminal repeat (LTR)-driven transcription of the RGR transfer plasmid produces a virus-like mRNA encoding the gRNA and containing a packaging signal sequence (Ψ, Fig 2B).

Modification of Gag with the FK506 binding protein FKB12 ensures effective packing of Gag/mRNA complexes following binding to the cell membrane. Addition of the rapalog (rapamycin analogue) ligand A/C during EV production mediates heterodimerisation of FRB (rapamycin-binding domain)-modified Cas9 with FKB12-Gag. The sgRNA encoded in virus-like mRNA is liberated in NanoMEDIC by self-cleavage of the flanking ribozymes and incorporated into *Streptococcus pyogenes* (Sp)Cas9. Dissociation of the complexes occurs after dilution of the A/C heterodimerizer once the EVs fuse with recipient cell membranes.

To test whether homology directed repair could be achieved following induction of double strand breaks, we generated a single-strand oligonucleotide (ssODN) template encoding a series of 6 stop codons interspaced by single nucleotides to allow for all possible open reading frames and ~50 nt homology arms (Fig 2A), as previously described [19]. The introduction of premature stop codons allows for degradation of modified transcripts by nonsense-mediated decay and/or premature termination of translation [20]. Overexpression of VSV-G in HEK293T cells leads to an increased production of VSV-G-expressing EVs, termed ‘gesicles’, that can be utilised for transfer of membrane, cytoplasmic and nuclear proteins [21]. We therefore loaded VSV-G-pseudotyped EVs with the ssODN template and tested whether it could be transferred to *N. brasiliensis* L3 (Fig 2D) [22,23].

### NanoMEDIC in conjunction with an extracellular vesicle-delivered homology directed repair template induces site directed mutagenesis in *Nippostrongylus* L3

In a first series of experiments, activated L3 were exposed to NanoMEDIC containing sgRNAs binding at nucleotide 46 and/or 91 of exon 3 in the presence or absence of EVs containing the STOP_ssODN for homology directed repair (HDR). Parasite genomic DNA was then assessed by PCR for modifications introduced by HDR (Fig 3A and B). PCR with primers binding the stop codon region and adjacent exon 2 or 4 led to amplification of expected PCR products (Fig 3C), indicating that NanoMEDIC in conjunction with EV-delivered ssODN templates could be used for site directed mutagenesis in *N. brasiliensis*. Furthermore, the presence of modified *dnase2* transcripts was confirmed by RT-PCR with an exon 2-specific forward primer and a reverse primer binding the 6xSTOP region, resulting in a product of approximately 140 bp (Fig 3D). PCR from genomic DNA with the same primer pair produced an amplicon of approximately 200 bp (Fig 3C). Importantly, the presence of modified cDNA provided evidence that NanoMEDIC and EV-delivered ssODN can reach tissues actively expressing DNase II following ingestion.

**Fig 3.**
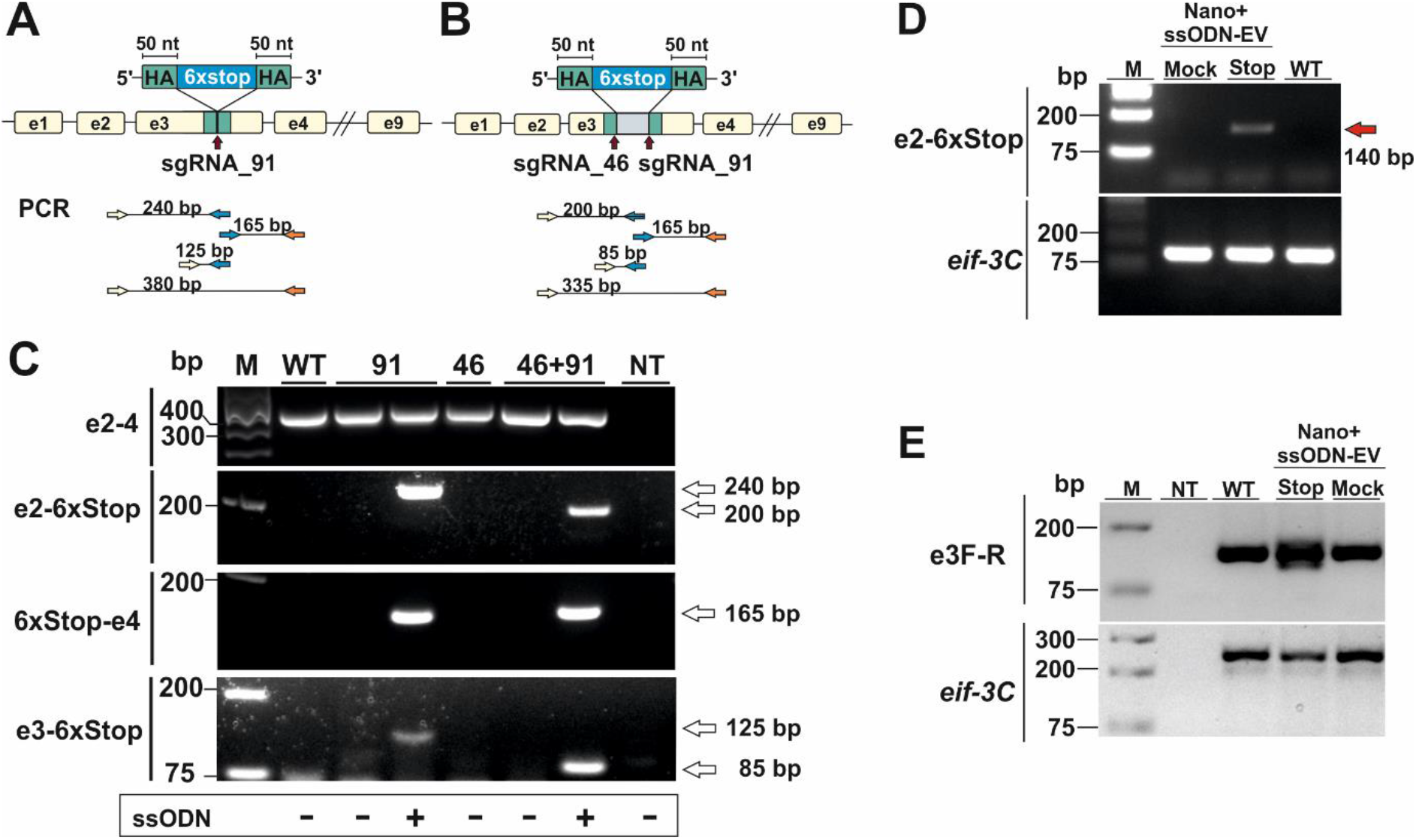
CRISPR/Cas9-mediated gene editing in *N. brasiliensis* infective larvae following delivery of Cas9/sgRNA complexes and homology directed repair templates via extracellular vesicles. (A,B) Knock-in of a homology directed repair (HDR) template expression cassette. HDR utilising a single stranded oligodeoxynucleotide (ssODN) with ~50 nt homology arms introduced a series of 6 stop codons following a single (B, sgRNA91) or double (C, sgRNA46 and 91) double stranded DNA break. (E) Detection of HDR sequences. PCR analysis of genomic DNA following exposure of activated L3 to NanoMEDIC with or without the addition of HDR-containing EVs. Primer pairs were designed to amplify fragments from the adjacent exon (2 or 4) and the HDR region (6xstop) and the respective amplicon sizes are indicated (A and B). Amplification of *eif-3c* (eukaryotic translation initiation factor 3 subunit C) was performed to control for gDNA integrity. WT, wildtype; NT, no template control. (D) Detection of modified *dnase2* transcript. Worms were exposed to EVs as above and modified *dnase2* transcripts confirmed by RT-PCR with an exon2-specific forward and HDR-specific reverse primer, resulting in a ~140 bp product. The same primer pair produces a 200 bp product from genomic DNA. (E) PCR of exon 3 with primers flanking the HDR integration site is indicative of insertions and deletions. Mock-Evs were electroporated with a single-strand oligonucleotide encoding an irrelevant sequence of the same length as the HDR template. *Eif-3C* was amplified to control for genomic DNA integrity. WT, wildtype; NT, no template control.

### Non-homologous end joining does not lead to effective mutations in L3

In previous studies with *Strongyloides stercoralis,* non-homologous end joining (NHEJ) of CRISPR/Cas9-induced double-strand breaks following microinjection of RNP complexes into the gonad resulted in large deletions, while smaller insertions or deletions (indels) appeared to be absent. Similarly, PCR of genomic DNA carried out with an exon 2-binding forward and an exon 4-binding reverse primer resulted in a single fragment of ~350 bp in all samples tested, which could not resolve possible indels (Fig 3C). To compensate for the possibility that deletions were insufficient, or frameshift not achievable using either guide, further experiments were carried out using a combination of both guide RNAs, with or without the addition of a ssODN allowing for deletion of the entire motif. To facilitate detection of smaller changes, a further PCR was carried out with primers flanking the double stranded break sites of exon 3 (Fig 3E). While multiple PCR fragments were generated in the NanoMEDIC + 6xSTOP_ssODN sample indicating integration of the HDR template, PCR with exon 3-binding forward and reverse primers still failed to resolve possible indels in worms exposed to NanoMEDIC (Fig 3E).

To examine potential low frequency errors induced by CRISPR/Cas9-mediated editing, we performed deep sequencing of PCR products derived from worms transfected with CRISPR/Cas9 either with or without the repair template. In worms transfected with CRISPR/Cas9 alone, after removing potential sequencing errors and PCR artefacts (see methods) single nucleotide polymorphisms (SNPs) at low frequency were detected mapping to the wild type allele of the DNase II gene, indicating that the enzyme could introduce breaks triggering error prone repair (Fig 4A). Interestingly, addition of the repair template led to an altered pattern of mutations in the wild type allele, without changing the frequency (Fig 4B). Similar patterns were seen for small indels mapping to the wild type allele (Fig 4C, D). The frequency of SNPs and indels mapping to the edited allele was low (Fig 4E,F), confirming that the homology directed repair was mostly accurate.

**Fig 4.**
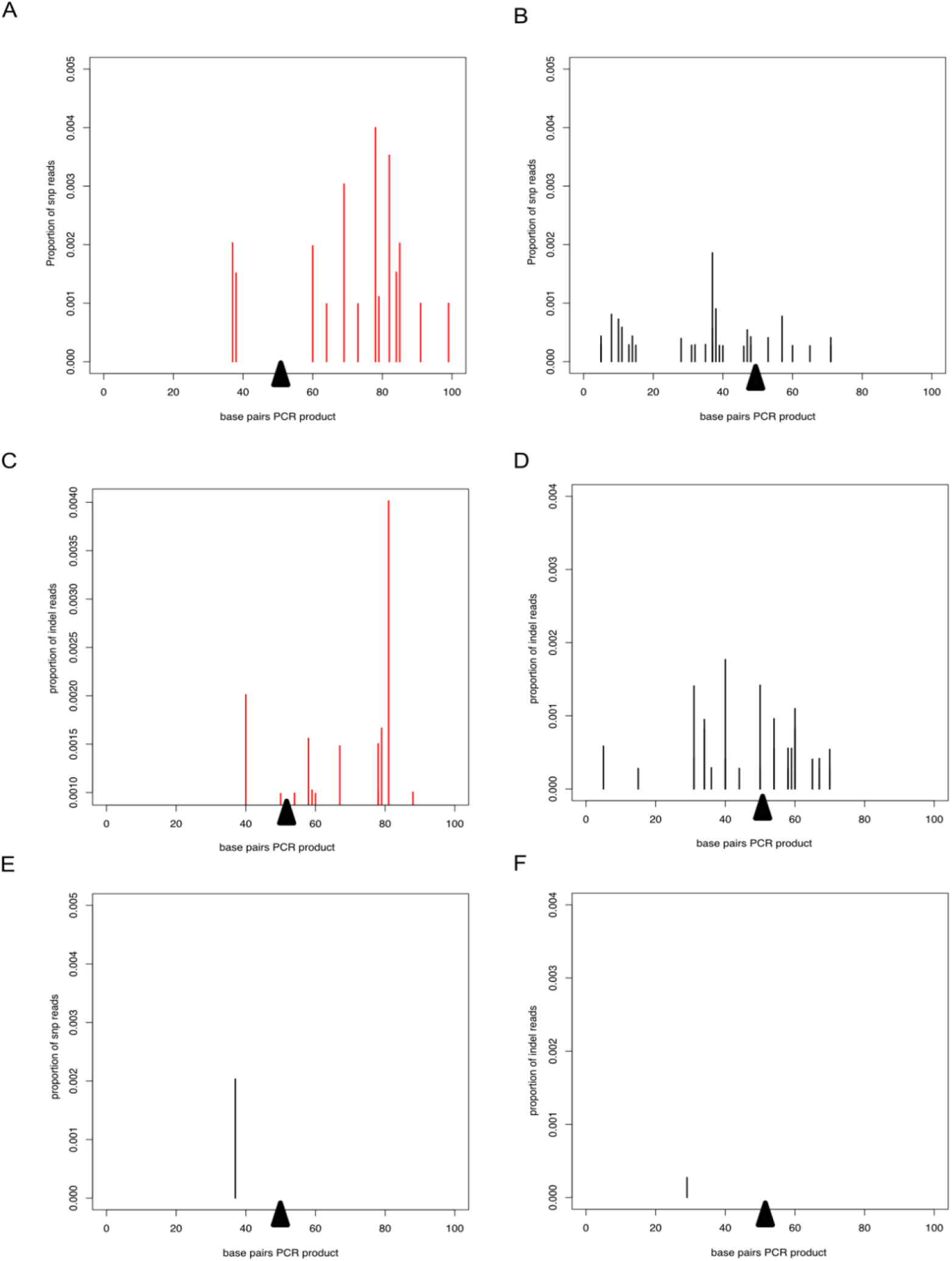
Deep sequencing of PCR product from worms transfected with CRISPR/Cas9 guide RNA complexes. In all plots, the proportion of reads corresponding to SNPs or indels is shown on the y axis and the presumed breakpoint is indicated by a black triangle. (A) and (B) SNPs on the WT allele after transfection with the CRISPR/Cas9 guide RNA complex without (A) or with (B) the repair template. (C) and (D) Indels on the WT allele for CRISPR/Cas9 guide RNA complex without (C) or with (D) the repair template. (E) SNPs and (F) indels mapping to the repair template after transfection with the CRISPR/Cas9 guide RNA complex and the repair template.

### Diminished DNase activity in CRISPR/Cas9 mutated larvae

We next assessed whether modifications introduced into the DNase II gene resulted in measurable reduction of secreted enzyme. Indeed, reduced DNase activity was observed in supernatants collected for 3 days following exposure of L3 to NanoMEDIC and ssODN (Fig 5A). While complete degradation of donor DNA was observed 5 min after exposure to supernatant from control worms, this required longer (10 min) incubation with supernatant of worms exposed to NanoMEDIC and the STOP_ssODN. Interestingly, delayed DNA degradation was only observed following delivery of the STOP_ssODN via VSV-G-EVs (ssODN-EV), while direct electroporation of NanoMEDIC with the STOP_ssODN (Nano+ssODN) was unsuccessful. Reduced nuclease activity was also not observed in supernatants from worms exposed to NanoMEDIC and Mock-EVs electroporated with a ssODN encoding an irrelevant sequence of the same length as the HDR template.

**Fig 5.**
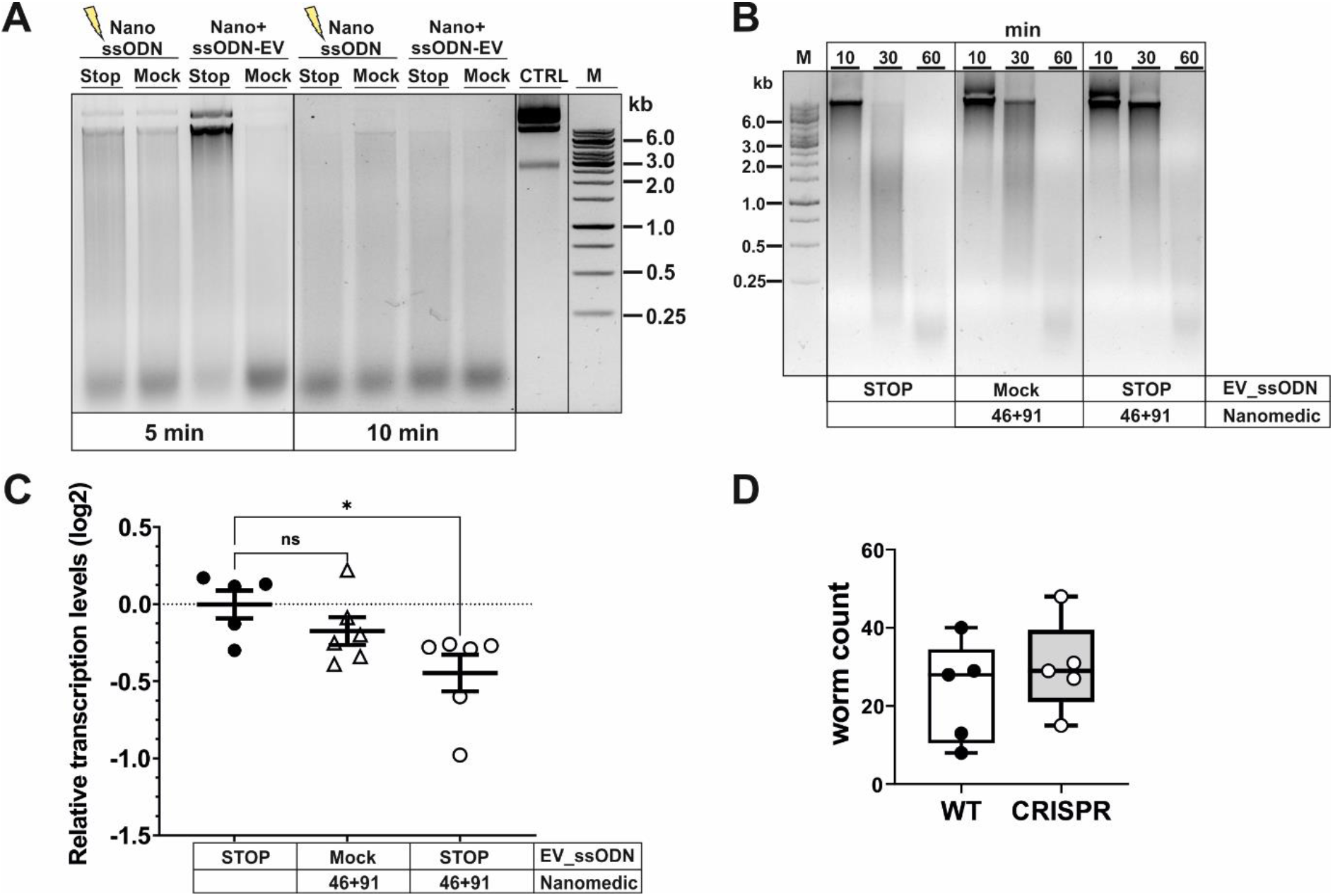
Editing of *dnase2* results in reduced enzyme activity in larval secreted products. (A) Reduced Dnase activity in larval secreted products. Activated L3s were exposed to Evs for 18 hours, washed and incubated for a further 48 hours. The HDR template was either electroporated into NanoMEDIC (Nano_ssODN) or VSV-G-pseudotyped EVs (ssODN-EV). Mock-EVs were electroporated with a ssODN encoding an irrelevant sequence of the same length as the HDR template. Larval secreted products were collected and assessed for DNase activity as described in Materials and methods. Undigested plasmid DNA (CTRL) is resolved on gels with test samples. (B) Time course of delayed DNA degradation by secreted products from modified worms. Activated L3s were exposed to EVs for 18 hours then incubated with plasmid DNA for 10, 30 or 60 min and supernatants analysed for DNA degradation by gel electrophoresis. (C) Downregulation of secreted *dnase2* transcripts in activated L3s after exposure to EVs. Transcript levels were assessed by RT-qPCR three days after transduction relative to wild type control larvae and normalised against the geometric mean of Ct values of reference genes *eif-3C* and *idhg-1.* Scatter plot with the mean ± sem of data from 2 independent experiments with 3 biological replicates each consisting of ~2,000 larvae. Treatment groups were analysed for significant differences with the Kruskal-Wallis test and Dunns *post-hoc* test in relation to the wild type. Statistical significance: p<0.05. (D) Reduction of DNase II secretion did not lead to lower parasite recovery in mice. Activated L3 were exposed to EVs for 18 hrs, washed and used to infect BALB/c mice. Adult worms were recovered from the intestines at day 5 post-infection and counted.

To assess the timeframe of delayed DNA degradation by modified worms, secreted products were assessed for their nuclease activity by adding a substrate plasmid DNA to the medium at the start of worm culture (Fig 5B). No intact DNA was detected after 30 min, and complete degradation recorded 60 min after culture with worms exposed to ssODN_EVs only. In contrast, co-culture of NanoMEDIC-exposed worms revealed the presence of some intact plasmid DNA after 30 min. This was more pronounced when worms were exposed to NanoMEDIC + ssODN_EVs, with some larger DNA fragments still present 60 min after co-culture (Fig 2B). Furthermore, RT-qPCR revealed a reduction in *dnase2* transcript levels by ~25% (mean log2 ± SEM = −0.45 ± 0.12) in NanoMEDIC + ssODN_EV samples, compared to the ssODN_EV only group (−0.003±0.09) (Fig 5C), indicating nonsense-mediated decay of modified transcripts. Some decrease of *dnase2* transcripts (−0.17±0.09) was also recorded in the NanoMEDIC-only group, but did not reach significance (p=0.74). These data, together with PCR analysis of genomic DNA, indicate that the majority of NanoMEDIC-induced gene disruption can be repaired by non-homologous end joining, and that editing is enhanced by homology-directed repair.

Because *N. brasiliensis* DNase II has been demonstrated to degrade neutrophil extracellular traps (NETs), it has been suggested to facilitate migration of L3 through the skin and lung tissues of their mammalian host [13]. However, the moderate silencing of the *dnase2* gene achieved in this study did not lead to a reduction in worm numbers in the intestines of infected mice (Fig 5D). Nevertheless, these data provide proof of principle that CRISPR/Cas9-induced gene editing can be achieved in infective larvae of *N. brasiliensis* by harnessing extracellular vesicle-mediated delivery of RNPs and HDR templates, providing a new route for genetic manipulation of parasitic nematodes.

### Expression of a NUC-1 orthologue by *Nippostrongylus brasiliensis*

As reduction of secreted DNase II activity did not result in altered recovery of adult worms from infected mice, we looked for additional enzymes in case this underlied some degree of redundancy in their action. A search of WormBase ParaSite revealed that NBR_0000088201 encoded an orthologue of lysosomal DNase II from *Caenorhabditis elegans* termed NUC-1 [24,25]. The derived amino acid sequence is shown in Fig 6A, aligned with that of the *N. brasiliensis* secreted DNase II (NBR_00001590) [13] and *C. elegans* NUC-1, revealing 65% identity between the mature *N. brasiliensis* and *C. elegans* NUC-1 proteins. Examination of transcript levels by real-time RT-PCR (qPCR) revealed that *Nb_nuc-1* is modestly upregulated in activated L3, consistent with emergence from a relatively quiescent state, then expressed at fairly constant levels through to adult worms (Fig 6B). Transcript levels for the secreted DNase II were 10-fold higher than *Nb_nuc-1* in activated L3, and at least 10-fold lower than *Nb_nuc-1* in resting L3 and adult worms (Fig 6C). This suggests that *Nb_nuc-1* has more of a housekeeping role, consistent with a lysosomal function, and it is notable that no nuclease activity was detected in culture supernatants of L3 within the first 2 hours following activation (Fig 1C) despite appreciable levels of *Nb_nuc-1* transcripts. In contrast, expression of the secreted DNase II was almost exclusively associated with larval activation, suggesting that this is the major or sole enzyme released into the mammalian host and primarily responsible for degradation of extracellular/environmental DNA in the early stages of infection.

**Fig 6.**
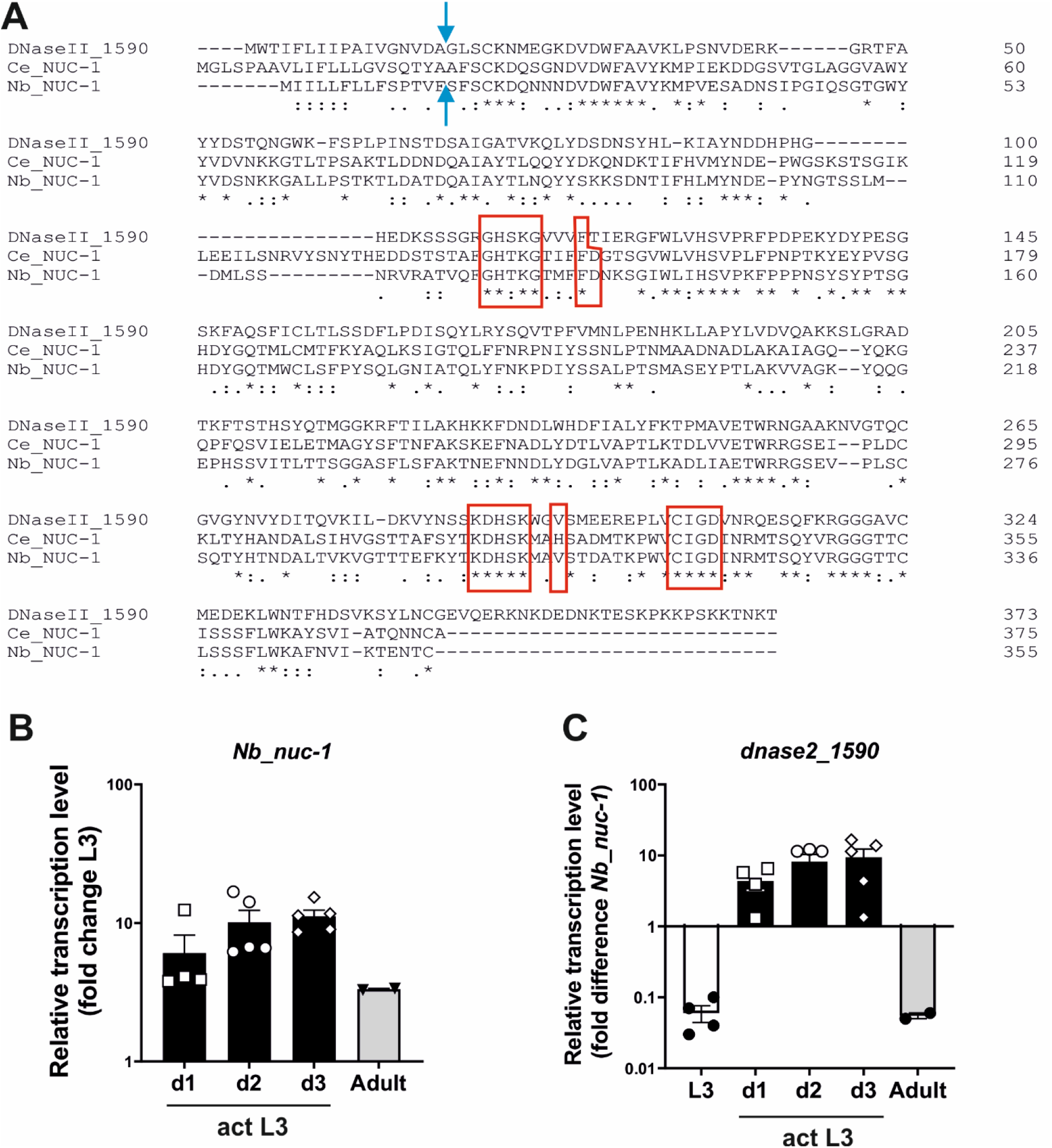
Detection of a *nuc-1* orthologue in *Nippostrongylus brasiliensis*. (A) Amino acid sequence of Nb_NUC-1 (NBR_0000088201) inferred by orthology to *C. elegans* NUC*-*1. Signal peptide cleavage site (blue arrow) and HKD motifs are indicated (red border). (B) *Nb_nuc-1* is upregulated in L3 following activation at 37°C. Transcript levels are shown relative to unactivated L3. (C) *Dnase2_1590* transcript levels relative to *Nb_nuc-1* in different life cycle stages and culture of L3 at 37°C up to 3 days. Interleaved scatter plot with bars and mean ± sem of 2 independent experiments with 2-3 biological replicates.

## Discussion

Nanoparticles, including extracellular vesicles (EVs), have become an important tool for the delivery of cargo including drugs, proteins, and nucleic acids to mammalian cells both in vitro and in vivo. EVs released by cells include microvesicles, exosomes, and apoptotic bodies, which differ in their biogenesis and size. However, while EVs have been described in secreted products of many helminths and suggested to be involved in signalling to the host [26], utilisation as vehicles to deliver functional cargo to parasitic helminths remains unexplored.

Here, we demonstrate that CRISPR/Cas9-mediated gene editing of *N. brasiliensis* is achievable using VSV-G-pseudotyped EVs (NanoMEDIC) for the delivery of RNP complexes. NanoMEDIC induced DNA double strand breaks, and homology directed repair was achieved through simultaneous delivery of a ssODN template via VSV-G-pseudotyped EVs (Fig. 1B). Edited transcripts were detected by RT-PCR, demonstrating that EVs can access cells of tissues actively expressing the gene of interest. Furthermore, gene disruption and the introduction of a repair template encoding 6 stop codons led to measurable reduction of target gene expression (*dnase2*), providing proof of principle that the technique can be employed for the identification and characterisation of molecules in parasites involved in disease processes.

As in previous studies with *Strongyloides stercoralis* [7], in the absence of a HDR template we could not detect smaller insertions or deletions following NHEJ. However, large genetic deletions of up to 500 bp in length following NHEJ have been described in *S. stercoralis* [7] and *Schistosoma mansoni* [26]. Due to the PCR design in our study, with primers spanning exon 2 to exon 4 of the DNase II gene, the presence of larger deletions cannot be excluded. However, detection of large deletions by whole genome sequencing may be challenging, as targeting tissues with direct access to EVs will result in mosaic genomes. The high abundance of wild type alleles masking edited genes has also been an issue in *S. stercoralis* [7]. While not reaching significance in our readouts so far, some reduction in transcript and DNase activity was observed in the absence of HDR, such that optimisation of NanoMEDIC production and purity may lead to improved gene disruption by NEHJ. However, consistent with previous studies in other species [7,19], gene disruption is more effective when HDR is employed, and it facilitates detection of edited genes by providing unique primer binding sites. Interestingly, while a ssODN template was sufficient to achieve HDR in *N. brasiliensis* L3 in this study, a double stranded DNA template was necessary for repair in *S. stercoralis* following injection of the gonad [7]. This may be indicative of different repair mechanisms in somatic and germline cells, as previously described in *C. elegans* [27].

Direct injection into the gonad has the advantage of potentially generating homozygous offspring, whereas NanoMEDIC targets tissues with direct access to EVs. The EVs used in this study are similar in size to lentiviral particles (~150-200 nm) and possess a cell membrane-derived envelope expressing VSV-G. Lentiviral particles can access cells in the intestine of *N. brasiliensis* L3, and interference with expression of secreted acetylcholinesterases suggests that they can also access subventral glands [5]. This is important, as it suggests that NanoMEDIC may be utilised for targeting tissues expressing secreted proteins, as shown here for the DNAse II. The same study showed that lentiviral particles can gain access to the germline, as integrated viral genomes were evident in a small proportion of the F1 generation [5]. The route to the germline is unclear, and further studies are required to determine whether this is similarly possible with NanoMEDIC.

Although we have shown that *N. brasiliensis* could be transduced with lentivirus, the transgene expression cassettes were subject to epigenetic silencing, and RNAi could not be maintained following development to adult stages [5]. CRISPR/Cas9-mediated gene editing offers a means to circumvent these problems, as more stable expression should be possible by site-directed integration of transgenes into regions less prone to epigenetic silencing. Lentiviral delivery of a Cas9 expression cassette has been superseded in *S. mansoni* by lipofection of in vitro assembled RNP complexes and simultaneous delivery of an HDR template by electroporation [19]. Furthermore, RNP complexes outperformed plasmid DNA-encoded Cas9 expression cassettes in *Strongyloides* [7]. Unlike *Strongyloides,* most parasitic nematodes do not have a free-living phase to facilitate microinjection. Using EVs for delivery of pre-assembled RNP complexes [12,29–31] and HDR templates [22] thus offers an alternative to techniques employed thus far, and pseudotyping EVs with VSVG allows for receptor-mediated uptake similar to viral transduction [12,29–31].

Due to their short half-life, RNPs are rapidly degraded, resulting in precise site-directed editing with low off-target frequencies [32]. Another major advantage is that RNPs are produced and assembled in mammalian cell lines for which optimised expression systems are readily available, avoiding lengthy optimisation of Cas9 expression cassettes for expression in the respective parasite. Use of mammalian cell lines facilitates cost effective, large scale production of EVs. Furthermore, in contrast to other vesicular delivery approaches relying on stochastic uptake of RNPs [29] or fusion of SpCas9 to Gag [27], NanoMEDIC utilise the Gag and FRB-FKB12 homing system to actively incorporate RNPs into budding EVs, and contain an average of seven Cas9 molecules per vesicle [12]. Moreover, VSV-G and Gag actively mediate the release of EVs from cells resulting in average titres in the range of 1 x 10^10^ particles per ml in our studies (S2 Fig).

Further optimisation of preparation and delivery of EVs will be necessary to improve editing efficiencies. Nanoflow analysis showed that NanoMEDIC preparations contained up to 90% exosomes (S2 Fig). Concentration in spin columns resulted in slightly less exosome (vesicles <100 nm) content than polymer-based precipitation (S2 Fig). NanoMEDIC have an average size of ~150 nm, and DNA is predominantly taken up by microvesicles (~150 nm) on electroporation [22]. The high content of exosomes is likely to saturate available receptors and impair effective uptake by NanoMEDIC. Polymer-based precipitation results in large quantities of lipids which may compete with available LDL receptors. Improved purity may thus require affinity chromatography [12] or a combination of filter columns [33].

In addition to loss of gene function by gene disruption, CRISPR/Cas9 provides a means to integrate foreign genes and expression cassettes via HDR [34]. Introduction of a traceable reporter or tag would allow sorting or enrichment of mutant larvae. It would also help to define the limitations of the EV delivery system in terms of which tissues can be accessed and manipulated, and facilitate investigation of gene expression patterns. For such studies, targeting a constitutively expressed, common gene such as tubulin, might be more effective.

Introduction of a reporter would require delivery of a dsDNA donor. Utilisation of a dsDNA template might allow editing in a wider range of tissues and improve HDR efficiencies as dsDNA templates usually have longer homology arms. Delivery of dsDNA via EVs is limited by their loading capacity, with an optimal length of DNA up to 750 bp but not exceeding 1000 bp [22], although loading capacities might be improved through optimisation of electroporation conditions [22,35].

In summary, we have demonstrated that EVs can be utilised as a vehicle to deliver functional cargo to a parasitic nematode and achieve CRIPSR/Cas9-mediated gene editing. Although the methodology clearly needs further development and optimisation, it should be applicable to a wide range of species, and could provide a new means for genetic manipulation of this important group of pathogens.

## Materials and Methods

### Ethics Statement

This study was approved by the Animal Welfare Ethical Review Board at Imperial College London, and was licensed by and performed under the UK Home Office Animals (Scientific Procedures) Act Personal Project Licence number PFA8EC7B7: ‘Immunity and immunomodulation in helminth infection’.

### Expression of recombinant DNase II in yeast

The coding sequence of *dnase2_1590* was amplified from *N. brasiliensis* cDNA omitting the signal peptide and stop codon and cloned into into pPICZalpha-A downstream of the coding sequence for the *Saccharomyces cerevisiae* α-mating secretion factor and in frame with an N-terminal myc and 6xHis Tag. PCR was carried out using Q5 polymerase (NEB) according to manufacturer’s recommendations with 500 nM of the following primers (lower case indicating nucleotides added for cloning purposes, restriction site underlined): F-5’-aagc<u>tGAATTC</u>GGTCTGAGTTGCAAGAACATGGAGG-3’ R-5-ttttg<u>tTCTAGA</u>GCGGTTTTGTTTGTCTTCTTGCTCG-3’

Following transformation of *Pichia pastoris* X-33, protein expression was optimised for single colonies and scaled up following the EasySelect Pichia expression protocol (Invitrogen). His-tagged proteins were purified from yeast supernatants by Ni-NTA resin affinity chromatography and protein concentration determined by Bradford assay.

### Antibody production and western blotting

A polyclonal antiserum to *N. brasiliensis* recombinant DNase II was raised by subcutaneous immunisation of a rat with 100 μg protein emulsified in alum, followed by 3 boosts of 50 μg protein via the same route at weeks 4, 6 and 8, and the animal bled at week 9. Western blotting was performed via standard procedures following resolution of 5 μg parasite proteins by SDS-12% polyacrylamide gel electrophoresis and blotting to polyvinylidene difluoride membrane. Rat anti-DNase II was used at 1:500 dilution, rabbit anti-rat IgG-horseradish peroxidase (Sigma) used as secondary antibody and the blot visualised using enhanced chemiluminescence western blotting detection reagents (Amersham Bioscience).

### RGR-vector construction

For facilitated cloning of synthesised sgRNA oligos, a NheI restriction site was introduced into the RGR expression vector (pL-5LTR-RGR(DMD#1)-AmCyan-A, Addgene plasmid #138482). To do this, the entire sequence between two SpeI restriction sites flanking the RGR region was amplified by PCR adding an NheI and AvrII restriction site to the 3’ end and used to replace the original sequence via SpeI and AvrII to allow for cloning of gRNA coding regions via KpnI and NheI. PCR was carried out in 50 μl reactions using Q5 polymerase (NEB) according to manufacturer’s recommendations with 1 ng of the RGR plasmid as template and 500 nM of the following primers (lower case indicating nucleotides added for cloning purposes and restriction site underlined): F-5’-GCTTGCATGCCGACATGGATTATTG<u>ACTAGT</u>CCC-3’; R-5’-attga<u>CCTAGGGCTAGC</u>TCTAGAGCGGCCGTCCCATTCGCCATGC-3.

The CRISPR/Cas-derived RNA-guided endonucleases (RGEN) algorithm was used to predict *Streptococcus pyogenes* (Sp) Cas9 gRNA targets with a 5’-NGG-3’ Protospacer Adjacent Motif (PAM) in exon 3 of the *dnase-2* gene and subsequent off-target screening against the *N. brasiliensis* genome. Oligonucleotides encoding gRNAs for subsequent integration into the RGR plasmid via KpnI and NheI were synthesized (GeneArt, Thermo Fisher Scientific) with the following structure: 5’-KpnI-inverted first 6 nt of the guide RNA-HH ribozyme-guide RNA-gRNA scaffold-HDV ribozyme-NheI-3’ (complete sequence Suppl 3). Plasmids were maintained in NEB stable *Escherichia coli* (NEB). Positive transformants were selected on LB agar plates containing 50 μg ml^-1^ ampicillin. Constructs were verified for error-free integration of transgenes by routine Sanger sequencing (Eurofins Genomics).

### Extracellular vesicle production

NanoMEDIC were produced in HEK293T cells maintained in Dulbecco’s Modified Eagle’s Medium (DMEM) at 37°C, 10% foetal calf serum (FCS), 2 mM L-glutamine, 100 units ml^-1^ penicillin and 100 μg ml^-1^ streptomycin, as described previously [12] with some modifications. In brief, per well of a 6-well plate, 3 x 10^6^ cells were transfected with 1.25 μg of gRNA-encoding plasmid, 1.25 μg of pHLS-EF1a-FRB-SpCas9-A (Addgene plasmid #138477), 1.25 μg of pHLS-EF1a-FKBP12-Gag^HIV^ (Addgene plasmid #138476), 250 ng of pcDNA1-Tat^HIV^ (Addgene plasmid #138478) and 500 ng of pMD2.G (VSV-G) (Addgene plasmid #12259), using Lipofectamine 2000 at a ratio of 1:2.5 (Life Technologies). After 16 hours, the transfection medium was replaced with 2 ml of reduced serum culture medium (5% FCS) supplemented with 300 μM A/C heterodimerization agent (formerly AP21967, Takara BioInc), 20 mM HEPES and 10 μM cholesterol (balanced with methyl-β-cyclodextrin, Sigma) [36]. VSV-G-EVs were produced in HEK293T cells following transfection of 3 x 10^6^ cells with 1.25 μg of pMD2.G per well of a 6-well plate with Lipofectamine 2000 at a ratio of 1:2.5. The cell culture supernatant was changed after 18 hours as for NanoMEDIC, omitting the heterodimerisation agent. After an additional incubation of 48 hours at 37°C and 10% CO_2_, the EV-containing cell supernatant was harvested, centrifuged at 2,000 x *g* for 20 min at 4°C and passed through a 0.45 μm Acrodisc syringe filter. NanoMEDIC and VSV-G-EVs were generally concentrated using 10 kDa vivaspin columns and washed twice with serum-free growth medium (NanoMEDIC) or trehalose buffer (10% trehalose in PBS) (VSV-G-EVs). Following centrifugation at 4,000 x *g* for 15 min at 4°C, the flow-through was discarded and EVs gently resuspended in 2 ml of the respective buffer before aliquoting into cryovials and storage at −80°C. For enrichment by polymer-based precipitation, 4 ml of Lenti-X concentrator (Takara Bio) was added to 12 ml of cell supernatant and the mixture incubated at 4°C with gentle agitation for 18 hours. Precipitated EVs were then pelleted by centrifugation at 3000 x *g* for 45 min and 4C and resuspended in 2 ml of trehalose buffer.

### Analysis of extracellular vesicles

The composition of EVs was analysed by Western blot and Nano-flow cytometry. Western blotting was performed via standard procedures following resolution of 12 μl concentrated EV preparations by SDS-12% polyacrylamide gel electrophoresis under reducing (Cas9, VSV-G) or non-reducing (CD63, CD81) conditions. Primary antibodies were used at 1:1000 dilution: mouse anti-human CD63 (clone H5C6, Biolegend); mouse anti-human CD81 (TAPA-1) (clone 5A6, Biolegend); rabbit anti-VSV-G (clone); mouse anti-CRISPR (Cas9) (clone 7A9, Biolegend).

Concentrated NanoMEDIC preparations were analysed for their nanoparticle content by nano-flow cytometry using a NanoAnalyzer calibrated against trehalose buffer. Before acquisition, samples were diluted 1:100 and 1:200 in trehalose buffer. The concentration of particles with diameters larger than 100 nm was determined following gating using the NanoFCM™ Silica Nanospheres Cocktail (S16M-Exo, diameter:68~155 nm) as a standard.

### Loading of VSV-G-EVs with HDR templates

Oligonucleotides encoding the HR templates (S3 Fig) were synthesized (Invitrogen) and reconstituted in nuclease-free water at a concentration of 1 μg μl^-1^. VSV-G-EVs were then loaded with ssDNA as described previously [22]. In brief, 5 μg of ssDNA were added to 95 μl of VSV-G-EVs in trehalose buffer and EVs then transferred to a 1 mm electroporation cuvette (BioRad) placed on ice. EVs were electroporated by exponential decay with two pulses at 400 V and 125 μF using a GenePulser Xcell electroporator (Bio-Rad). Cuvettes were placed on ice immediately after electroporation and incubated on ice for 20 min. EVs were then transferred to fresh microfuge tubes and the cuvette washed with one volume (100 μl) of RPMI and added to the tube. To alleviate aggregation, EDTA was added to a final concentration of 1 mM and EVs incubated at room temperature for 15 min, gently resuspended several times during incubation [22].

### Parasite infection, recovery and exposure to EVs

*N. brasiliensis* were maintained in male SD rats, and infective larvae isolated from faecal cultures using a Baermann apparatus. Larvae were activated to feed as previously described [15] for 48 to 72 hrs in RPMI1640, 0.65% glucose, 2 mM L-glutamine, 100 U ml^-1^ penicillin, 100 μg ml^-1^ streptomycin, 100 μg ml^-1^ gentamicin, 20 mM HEPES, 2% rat serum (worm culture medium), then washed twice in serum-free medium prior to exposure to EVs. Per well of a 12-well plate, approximately 3,000 - 4,000 activated L3 were exposed to 200 μl NanoMEDIC and/or 200 μl of VSV-G-EVs, volumes adjusted to 1 ml with serum-free medium and 10 μg ml^-1^ polybrene (Sigma) and 200 μg ml^-1^ gentamicin added. EV preparations in control worms were substituted with HEK293T cell supernatant from untransfected cells. Following incubation for 18 to 24 hrs, worms were transferred to a 15 ml tube and washed twice in 10 ml of warm serum-free worm medium containing 1 mM EDTA with centrifugation at 150 x *g* for 1 min between washes. Worms were then resuspended in 2 ml complete worm medium (or serum-free medium when testing for ES products) and incubated for another 24 to 48 hours at 37°C, 5% CO_2_.

### Detection of HDR in genomic DNA

For integration studies in L3s, worms were washed twice in 10 ml of warm PBS at 72 hrs post exposure to EVs and genomic DNA isolated using the DNeasy Blood and Tissue DNA extraction kit (Qiagen). Modified DNA was detected by 2-step PCR with an annealing/extension temperature of 72°C using Q5 polymerase and the following primers (F: forward; R: reverse): e2F: 5’-GATTCGGCTATTGGTGCAACTGTTAAGC-3’; e3F: 5’-ACCTCAAAATTGCCTACAACGAC-3’; e3R: 5’-GGAATCTTGGCACACTGTGTACCAGC-3’; e4R: 5’-TCGAGCCTGATTCGGGGTAGTCG-3’; StopF: 5’-TAAGTGACTAGGTAACTGAGTAGC-3’; StopR: 5’-ATCCCCGTGCTACTCAGTTACCTAGTCACTTA-3’.

### Preparation of DNA libraries and deep sequencing

DNA libraries for deep sequencing were made using the NEBNext Ultra II DNA library preparation kit (NEB) according to the manufacturer’s instructions. Sequencing was performed by the London Institute of Medical Sciences Genomics facility. Reads were mapped to the predicted PCR products from wild type and edited alleles using Bowtie2 [37]. SNPs and indels were mapped using samtools mPileup [38]. We then used Varscan2 [39] using the commands pileup2snp and pileup2indel to identify putative SNPs and indels respectively, requiring a minimum average quality of 20 and setting a minimum frequency of 1 in 10000 to enable low frequency alterations to be detected. Data were read into R using the read.table function and comparison of the different files enabled called SNPs that were present in the untransfected control to be removed as likely sequencing errors or PCR artefacts. Line plots indicating the frequency of SNPs at each position divided by total reads mapping to that position were constructed to illustrate the distribution of mutations along the template.

### Reverse Transcription PCR (RT-PCR) and Real-Time quantitative PCR (RT-qPCR)

Total RNA was extracted using TRIreagent (Sigma) and converted to cDNA using an iScript cDNA kit (Biorad) following removal of contaminating genomic DNA by DNAse I. RT-PCR was carried out using Q5 DNA polymerase (NEB) according to the manufacturer’s recommendations. RT-qPCR was carried out with a Step-One PLUS Fast Real-time PCR cycler (Applied Biosystems) under standard fast cycling conditions using PowerUP SYBR Green PCR Master Mix (Applied Biosystems) and 250 nM target gene specific forward and reverse primers. PCR amplification efficiencies were established for each primer pair [40] and ranged between 1.9 and 2.1. Cycle threshold (Ct) values of target genes were normalised to the geometric mean of *eif-3C* (NBR_0001150401) and *idhg-1* (NBR_0000658601) [5] and calibrated to the mean untreated control (wild type) samples for relative quantification by the comparative Ct method [41]. The primers were (forward (F), reverse *(R)):Nb-nuc-1* F: 5’-TGACGAACCATACAACGGCA-3’, R: 5’-TGGAACACTGTGGATCAGCC-3’; *eif-3C* F: 5’-GAACACGTTGTAGCTGCGTCA-3’, R: 5’-AATAGGTTCTCAGCGATTCCGTT-3’; *idhg-1* F: 5’-CAGAAATTGGGAGACGGCCT-3’, R: 5’-CCGAGAAACCAGCTGCATAGA-3’; dnase2_1590_E6F: 5’-TGGAAACTTGGAGAAACGGTGCTG-3’; dnase2_1590_E7R: 5’-ACATCTCCGATACAAACTAGGGGCTCC-3’; e2F: 5’-GATTCGGCTATTGGTGCAACTGTTAAGC-3’; StopR: 5’-ATCCCCGTGCTACTCAGTTACCTAGTCACTTA-3’.

### DNase activity assay

Supernatant collected from worms cultured in serum-free medium was thawed and all reactions prepared on ice. Per test sample, 500 ng of a plasmid DNA substrate in 10 μl nuclease free water was placed in PCR tubes and a 10 μl droplet of worm supernatant transferred to the side of the tube. Reactions were initiated by centrifugation and samples placed immediately into a PCR cycler pre-warmed to 37°C and incubated for 2 to 10 min. After heat inactivation at 75°C for 10 min, 4 μl of agarose loading dye was added and 10 μl of the sample separated on a 1% agarose gel.

To assess the dynamics of DNase secretion by L3, worms were extensively washed in serum-free medium, counted and resuspended at 2500 L3 ml^-1^. Per time point tested, 80 μl of worm suspension (200 L3) was added to 20 μl of plasmid DNA solution (2 μg in 20 μl RPMI).

Following incubation for 15, 30, 60 or 120 min at 37°C, 5% CO_2_, 80 μl of supernatant was carefully aspirated and transferred to a PCR tube. Samples were immediately frozen at −20°C. For analysis, tubes were placed directly from the freezer into a PCR cycler prewarmed to 75°C and heat inactivated for 10 min. Loading dye was added and the samples separated on a 1 % agarose gel.

## Statistics

Treatment groups were analysed for significant differences with the Kruskal-Wallis test and Dunns *post-hoc* test in relation to the control group. Values are expressed as the median with range or the mean ± SEM, and significant differences were determined using GraphPad Prism. P values of <0.05 were considered significant, *p<0.05.

## Author Contributions

Conceived and designed the experiments: JH MES PS. Performed the experiments: JH SG MES PS. Analyzed the data: JH PS MES. Wrote the paper: JH MES PS.

## Acknowledgements

This study was funded by a BBSRC grant to MES, PS and JH (BB/S001085/1).

## Supporting Information

**S1 Fig.**
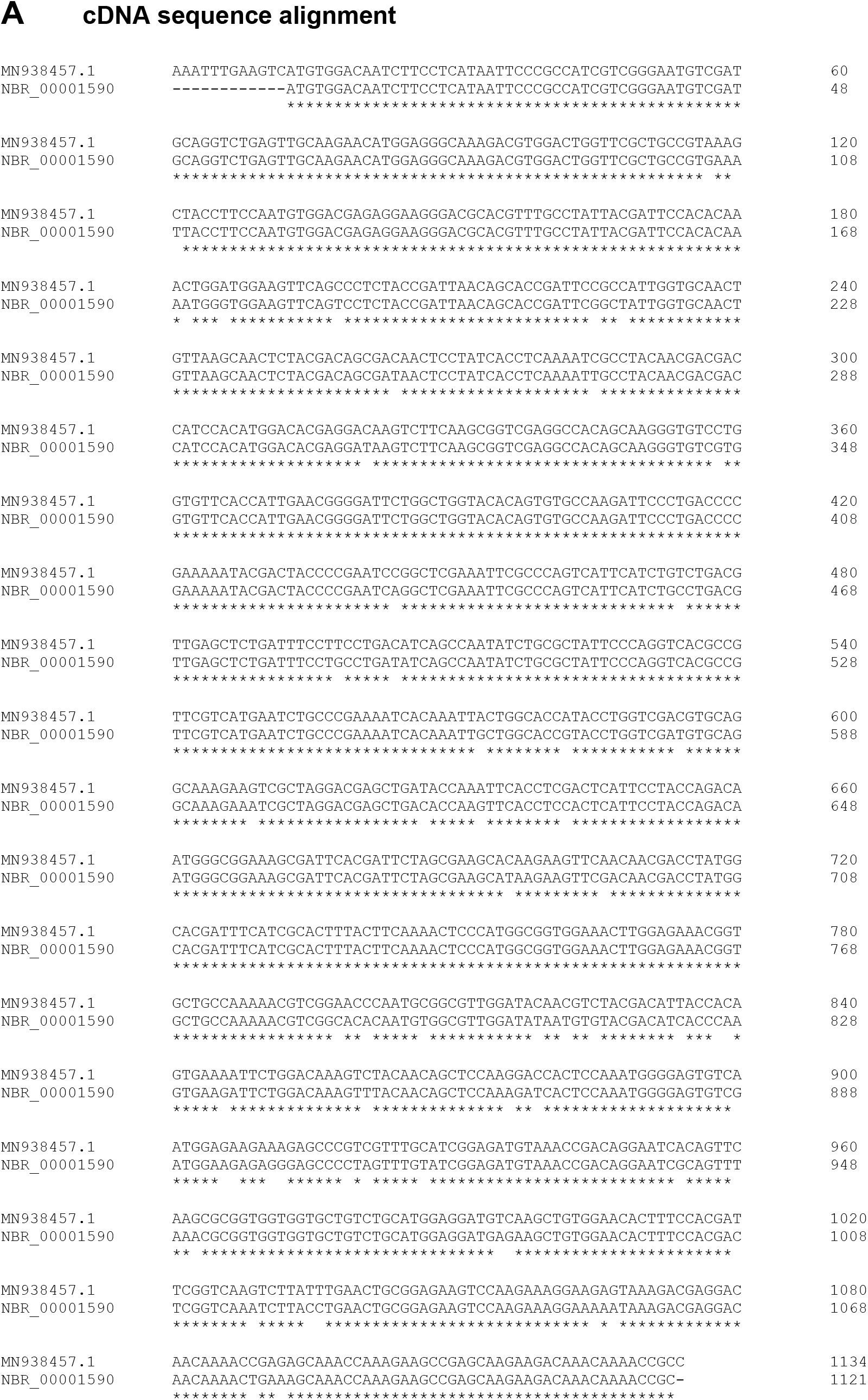
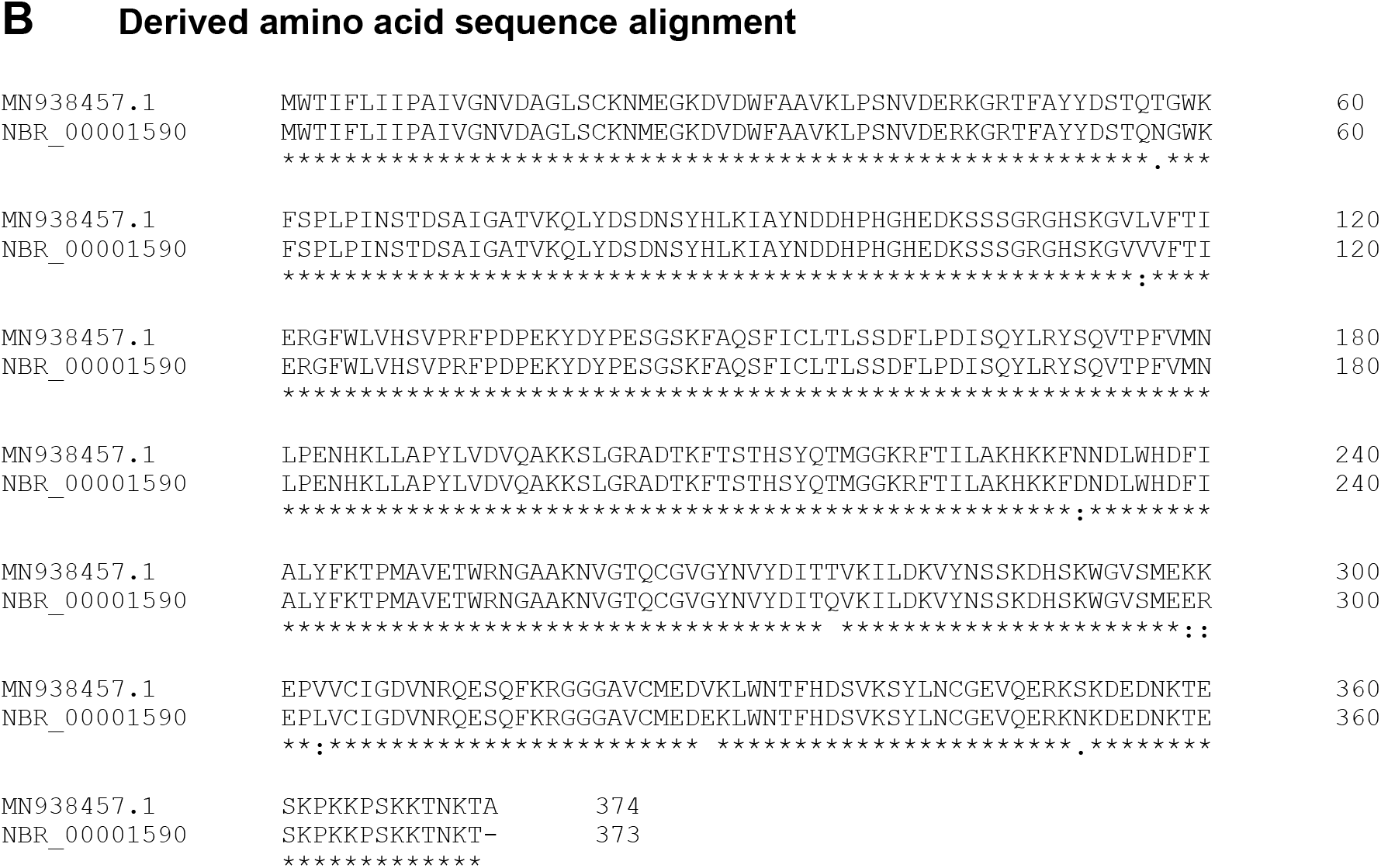
Alignment of cDNA and derived amino acid sequences for MN938457.1 (GenBank) and NBR_00001590 (Wormbase ParaSite).

**S2 Fig. Analysis of extracellular vesicle preparations**

### A Analysis of extracellular vesicle production by western blot

Extracellular vesicles (EV) or NanoMEDIC (Nano)-containing cell supernatants were concentrated using vivaspin columns (VIVA) or precipitation with Lenti-X concentrator (LX) as described in Materials and methods. Western blotting was performed following SDS-PAGE under reducing conditions to determine the presence of Cas9 and VSV-G, or under non-reducing conditions for the presence of CD63 and CD81.

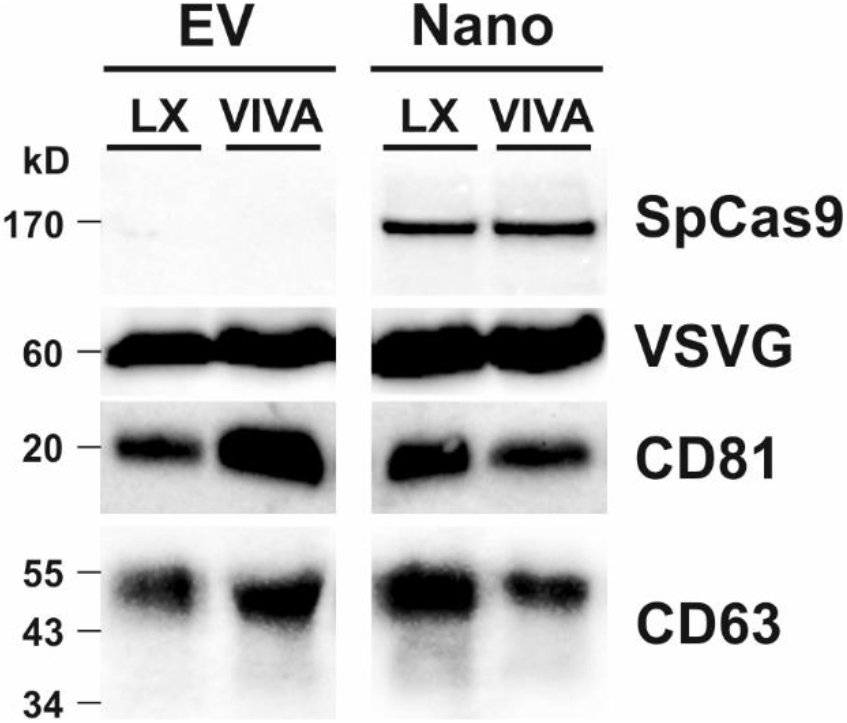

### B Analysis by Nano-flow cytometry

NanoMEDIC preparations concentrated by ultrafiltration (VIVAspin) or precipitation (Lenti-X) were analysed for their nanoparticle content by nano-flow cytometry as described in Materials and methods. The table shows the concentration and proportion of particles with diameters larger than 100 nm.

|  | *Lenti-X* | *VIVAspin* |
| --- | --- | --- |
| *Sample concentration* | $1.88 \times 10^{11}/\text{ml}$ | $5.12 \times 10^{10}/\text{ml}$ |
| *Gated EVs (>100 nm)* | 7.7% | 16.19 % |
| *Gated EVs (>100 nm) concentration* | $1.45 \times 10^{10}/\text{ml}$ | $0.83 \times 10^{10}/\text{ml}$ |

**S3 Fig.**
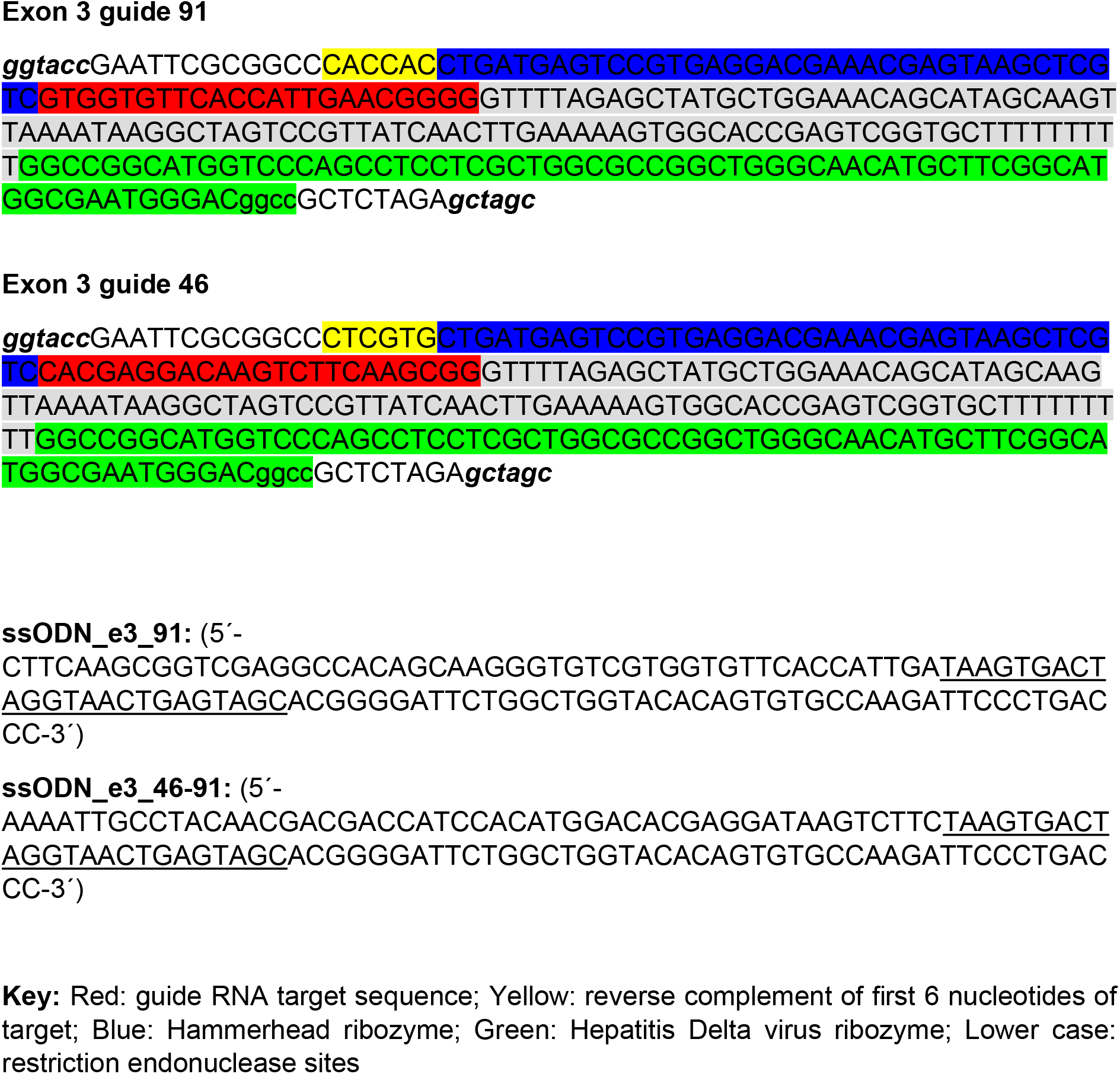
Guide RNA-encoding region synthesised for cloning into RGR plasmid and ssODN sequences for homology directed repair.

